# Limiting Pool and Actin Architecture Controls Myosin Cluster Sizes in Adherent Cells

**DOI:** 10.1101/2023.06.07.544121

**Authors:** Wen-hung Chou, Mehdi Molaei, Huini Wu, Patrick W. Oakes, Jordan R. Beach, Margaret L. Gardel

## Abstract

The actomyosin cytoskeleton generates mechanical forces that power important cellular processes, such as cell migration, cell division, and mechanosensing. Actomyosin self-assembles into contractile networks and bundles that underlie force generation and transmission in cells. A central step is the assembly of the myosin II filament from myosin monomers, regulation of which has been extensively studied. However, myosin filaments are almost always found as clusters within the cell cortex. While recent studies characterized cluster nucleation dynamics at the cell periphery, how myosin clusters grow on stress fibers remains poorly characterized. Here, we utilize a U2OS osteosarcoma cell line with endogenously tagged myosin II to measure the myosin cluster size distribution in the lamella of adherent cells. We find that myosin clusters can grow with Rho-kinase (ROCK) activity alone in the absence of myosin motor activity. Time-lapse imaging reveals that myosin clusters grow via increased myosin association to existing clusters, which is potentiated by ROCK-dependent myosin filament assembly. Enabling myosin motor activity allows further myosin cluster growth through myosin association that is dependent on F-actin architecture. Using a toy model, we show that myosin self-affinity is sufficient to recapitulate the experimentally observed myosin cluster size distribution, and that myosin cluster sizes are determined by the pool of myosin available for cluster growth. Together, our findings provide new insights into the regulation of myosin cluster sizes within the lamellar actomyosin cytoskeleton.

## Introduction

Various cellular processes and dynamics in cells depend on mechanical forces generated by the actomyosin cytoskeleton, such as cell motility (Vicente-Manzanares *et al*., 2009), adhesion (Oakes *et al*., 2012; Stricker *et al*., 2013), and mechanosensing (Lee and Kumar, 2016). The central contractile molecular element of the actomyosin cytoskeleton is non-muscle myosin II (NMII, hereafter referred to as myosin), which self-assembles into myosin filaments that contract actin filaments (F-actin) to generate contractile forces. In combination with crosslinkers and other actin binding proteins, actin and myosin build contractile actomyosin structures that underlie force production in cells. For example, in non-muscle cells, F-actin bundles and myosin filaments form contractile bundles that span the cell, called stress fibers. While both F-actin and myosin must be regulated to build stress fibers, F-actin regulation has been well-studied (for example, reviewed in Pollard, 2007; Kadzik *et al*., 2020) and myosin regulation is relatively less understood.

Most studies on myosin regulation have focused on the assembly of myosin filaments from myosin molecules. At the molecular scale, the ability of myosin to assemble into myosin filaments is determined by the transition between its assembly-incompetent autoinhibited state and its assembly-competent active state (Quintanilla *et al*., 2023b). This transition largely depends on phosphorylation on the regulatory light chain (RLC) of myosin. Several kinases, including the myosin light chain kinase (MLCK) and Rho-associated coiled-coil containing kinase (ROCK), can phosphorylate the Thr18 or Ser19 residues of the RLC to promote myosin filament assembly (Scholey *et al*., 1980; Vicente-Manzanares *et al*., 2009; Newell-Litwa *et al*., 2015). Phosphorylation sites on the myosin heavy chain has also been shown to play a role (Vicente-Manzanares *et al*., 2009; Dulyaninova and Bresnick, 2013). Aside from biochemical regulation, mechanical tension has also been proposed to impact the assembly of myosin filaments (Luo *et al*., 2012; Shutova *et al*., 2012; Grewe and Schwarz, 2020).

Despite the importance of myosin filament regulation, single myosin filaments are rarely observed in cells. While some individual myosin filaments may be observed near the cell periphery at the leading edge, most myosin filaments in cells form clusters, where multiple myosin filaments come into close contact with each other with their myosin head domains. Described as ribbons or stacks, myosin clusters were best visualized using electron microscopy (Verkhovsky *et al*., 1995; Svitkina *et al*., 1997; Shutova *et al*., 2012, 2014). Myosin clusters at the leading edge of the cell usually contain fewer myosin filaments, while more myosin filaments cluster together towards the cell body as they associate with stress fibers. Myosin clusters can further register with each other to form super-structures on stress fibers (Hu *et al*., 2017). However, it remains largely unknown how myosin filaments self-associate to form clusters and how the process is regulated.

Recent studies have characterized the dynamics of myosin clusters to understand their formation at the leading cell edge. Using super-resolution microscopy, myosin filaments were observed nucleating at the leading edge of the cell (Fenix *et al*., 2016; Beach *et al*., 2017). After nucleation, more myosin filaments were recruited to the nucleated filament to form myosin clusters. Through rounds of amplification and splitting, myosin clusters amplify both in size and number before incorporating into stress fibers. The amplification and splitting dynamics have been shown to depend on myosin monomer availability, myosin motor activity, actin dynamics, and actin density (Fenix *et al*., 2016; Beach *et al*., 2017). While myosin cluster dynamics at the leading edge and their regulation is thought to be general for all myosin clusters in cells, myosin cluster sizes on stress fibers have not been properly characterized.

In this work, we explore the regulation of myosin cluster sizes distribution in the lamella of adherent cells. To this end, we quantitatively characterized endogenously tagged non-muscle myosin IIA (NMIIA) in adherent U2OS osteosarcoma cells. The size distribution of myosin clusters found within actin networks and bundles within the lamella is quite broad. ROCK activity is sufficient to grow myosin clusters without myosin motor activity by increasing the pool of assembled myosin that can associate with existing ROCK-independent myosin clusters. Myosin motor activity further enhances myosin cluster growth by myosin association that is dependent on F-actin architecture. A toy model of myosin cluster growth with myosin self-affinity is sufficient to recapitulate the broad distribution of cluster sizes. Our results suggest that myosin cluster sizes within the lamella of adherent cells are set by a limiting pool of myosin available for cluster growth.

## Results

### Myosin filaments form clusters of a broad range of sizes within lamellar actin networks and bundles

To study the regulation of myosin clusters, we chose the human osteosarcoma U2OS cell line because their lamellar actin architectures, including stress fibers, have been well-characterized (Hotulainen and Lappalainen, 2006; Tojkander *et al*., 2012). Using a CRISPR knock-in approach, we endogenously labeled non-muscle myosin IIA by inserting an mScarlet gene at the C-terminal locus of MYH9 (see methods). We chose to focus on non-muscle myosin IIA because of its role in generating contractile forces in cells (Chang and Kumar, 2015; Weißenbruch *et al*., 2021), and it is the dominant myosin isoform in U2OS cells. Since myosin assembles into a filament with their C-terminal tail overlapping at the bare zone, the fluorescence signal represents the central location of a myosin filament (Fig. 1A). Using spinning disk confocal imaging, we see that most myosin appears as diffraction-limited puncta of similar physical dimensions but with varying intensities (Fig. 1B). These myosin puncta often colocalize with F-actin bundles (Fig. 1B). Since our imaging conditions are likely insufficient to confidently detect single myosin filaments, each punctum likely contains multiple myosin filaments. We therefore refer to these punctate structures as myosin clusters, which may represent single or multiple closely associated myosin stacks.

**Figure 1.**
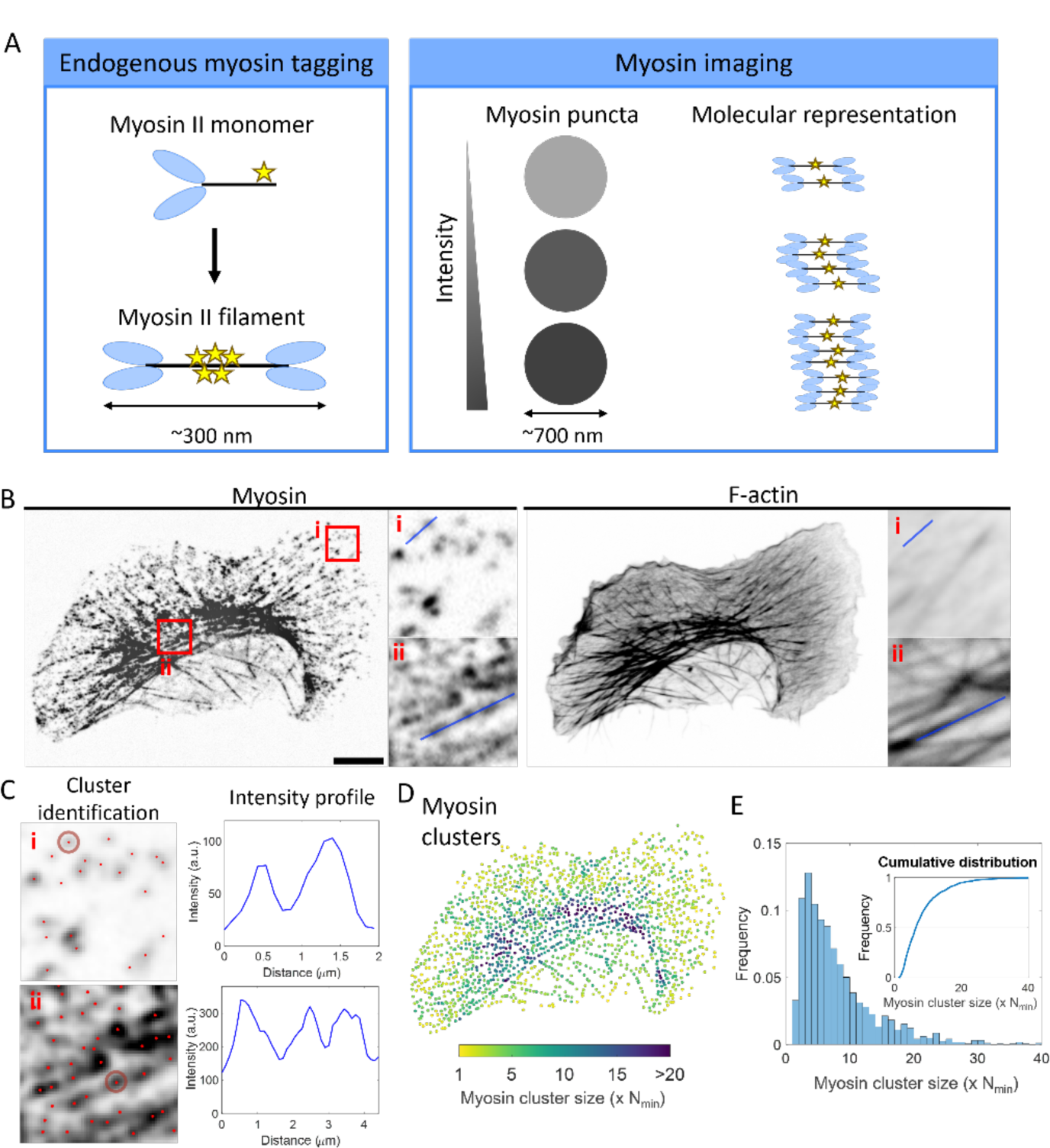
Myosin filaments form clusters of a broad range of sizes on stress fibers. (A) Schematics of the experiment setup. Non-muscle myosin IIA is endogenously tagged on the C-terminal tail so that myosin filaments will appear as diffraction-limited puncta. The intensity of each punctum is directly proportional with the amount of myosin filaments in each cluster. The number of myosin heads on a filament have been reduced for simplicity. (B) Representative images of non-muscle myosin IIA (myosin) and F-actin in a U2OS cell. The right panels show zoom-in of myosin clusters from the two red boxes. Images are inverted. Scale bar 10 µm. (C) Examples of myosin cluster localization. In the cluster identification column, each myosin intensity maxima is shown as a red dot. The brown circles indicate a typical area where the integrated intensity of a myosin punctum is calculated. The intensity profile column shows the intensity profile along the blue line in (B)i and (B)ii. (D) Myosin clusters in cells color-coded by their sizes. N_min_ represents the minimal myosin cluster size that can be detected, as discussed in the text. (E) Histogram of myosin cluster sizes. Inset shows the cumulative distribution of myosin cluster sizes.

To characterize myosin cluster dynamics, we conducted time-lapse live-cell imaging. While myosin clusters undergo retrograde motion and contract inward towards the cell center, they remain stable over time, persisting for at least 8 minutes (Supp. Fig. 1A & Supp. Movie 1). This suggests that myosin clusters are not multiple randomly overlapping structures but rather stable structures that move together as one unit. Despite their structural stability, the constituents of myosin clusters exchange rapidly. When we perform fluorescence recovery after photobleaching (FRAP) analysis on myosin clusters, we see that myosin fluorescence recovers with a half-time of ∼2 minutes and a mobile fraction of 80% (Supp. Fig. 1B, D, E), consistent with previous reports (Hotulainen and Lappalainen, 2006; Breckenridge *et al*., 2009; Chang and Kumar, 2015; Weißenbruch *et al*., 2022). We also observed FRAP recovery at the level of individual myosin clusters. Instead of assembling new myosin clusters or transporting clusters into the photobleached region, nearby myosin clusters undergo FRAP recovery at a similar rate (Supp. Fig. 1C). This suggests that myosin is continually exchanging within myosin clusters, with constant association and dissociation of myosin. Since myosin clusters contain multiple myosin filaments (see characterization below), FRAP recovery may indicate the exchange of single myosin monomers, oligomers, or whole filaments.

Since myosin is endogenously labeled, myosin punctum intensity reflects the number of myosin filaments within each cluster. To quantify the intensities of myosin clusters bound to the lamellar actin network or bundles, we fixed and permeabilized the cells before imaging to remove unbound or weakly-bound myosin. We noticed diffuse myosin intensity within the cell that is higher than background intensities even after fixation and permeabilization (Supp. Fig. 2); this may indicate myosin bound to the cellular cortex. To focus on quantifying the intensities of myosin puncta, we exclude the diffuse background by performing local background subtraction. Then, the centroid location of each myosin punctum are determined using previously developed methods (Crocker and Grier, 1996). Briefly, each image is passed through a bandpass filter to preserve circular features with a diameter of ∼700 nm, about the typical size of a myosin punctum. Local maxima with intensity higher than background noise are accepted as a puncta candidate (Fig. 1C). We further filter out puncta that are irregularly shaped or puncta whose intensity are too spread out (see methods). The remaining puncta candidates are accepted as valid myosin clusters. Each myosin cluster’s intensity is then calculated as the integrated intensity of each punctum within a diameter of ∼700 nm.

While the absolute number of myosin molecules within each cluster requires careful calibration between fluorescence intensity and the number of fluorescent proteins (Quintanilla *et al*., 2023a), their relative sizes can be assessed without this calibration. To compare the sizes of myosin clusters across different cells and experimental conditions, we normalize all myosin cluster intensities within the same set of experiment against the minimum myosin intensity identified in each experiment. We posit that this minimum intensity (I_min_) reflects the minimal number of myosin molecules in a cluster (N_min_) we can detect above the noise floor. N_min_ likely represents more than one myosin filament (see Quintanilla *et al*., 2023, which suggests that the vast majority of identifiable myosin structures at the leading edge of the cell contains more than one myosin filaments). Since we keep the imaging conditions and analysis parameters constant in each experiment, we reasoned that N_min_ will be the same but I_min_ can vary slightly across different experiments. Therefore, we normalize myosin cluster intensities in each experiment to I_min_ and present myosin cluster sizes as multiples of N_min_.

With this approach, we find a broad distribution of myosin cluster sizes across the cell, ranging up to 30-fold. Most myosin clusters contain 3 times more myosin than N_min_, but there is a sizeable fraction of larger myosin clusters containing 20-fold more myosin than N_min_ (Fig. 1D, E). This is also evidenced by the histogram of myosin cluster intensities, which shows a long-tailed distribution (Fig. 1E). Myosin intensity is also spatially heterogeneous (Fig. 1D). Myosin clusters tend to be smaller at the lamellar region or newly formed transverse arcs, containing only 2-4 times more myosin than N_min_ (Fig. 1C, i). On the other hand, myosin clusters tend to be larger on mature transverse arcs in the cell body or on ventral stress fibers, with more than 15 times more myosin than N_min_ (Fig. 1C, ii).

### Rho-kinase activity drives growth through net myosin association to existing clusters

Recent literature showed that myosin clusters at the leading edge is regulated by myosin monomer availability and motor activity (Fenix *et al*., 2016; Beach *et al*., 2017). We designed an experiment to test their respective contributions to the growth of myosin clusters within the lamellar actin. To promote myosin cluster disassembly, cells were treated with 40 µM ROCK inhibitor Y-27632 (Y-27, inhibits ROCK-dependent myosin light chain phosphorylation) and 50 µM blebbistatin (myosin II ATPase inhibitor and locks myosin in a weak actin-binding state (Kovács *et al*., 2004)) for 30 minutes. Even after treatment, residual myosin clusters with low fluorescence intensity remained (Fig. 2A, Y+B+). Compared with the cluster size distribution of control cells, the peak was shifted towards smaller cluster sizes, and the upper tail of the distribution was reduced (Fig. 2B). This indicates that the sizes of the remaining myosin clusters are smaller with less variation, averaging around 2-3 times the size of N_min_ (Fig. 2B, Y+B+). These myosin clusters are presumably formed by myosin filaments that are phosphorylated by other kinases, such as MLCK, MRCK, or CDC42, or formed by increasing the total monomer pool above a critical threshold to enable spontaneous filament assembly (Quintanilla *et al*., 2023a).

**Figure 2.**
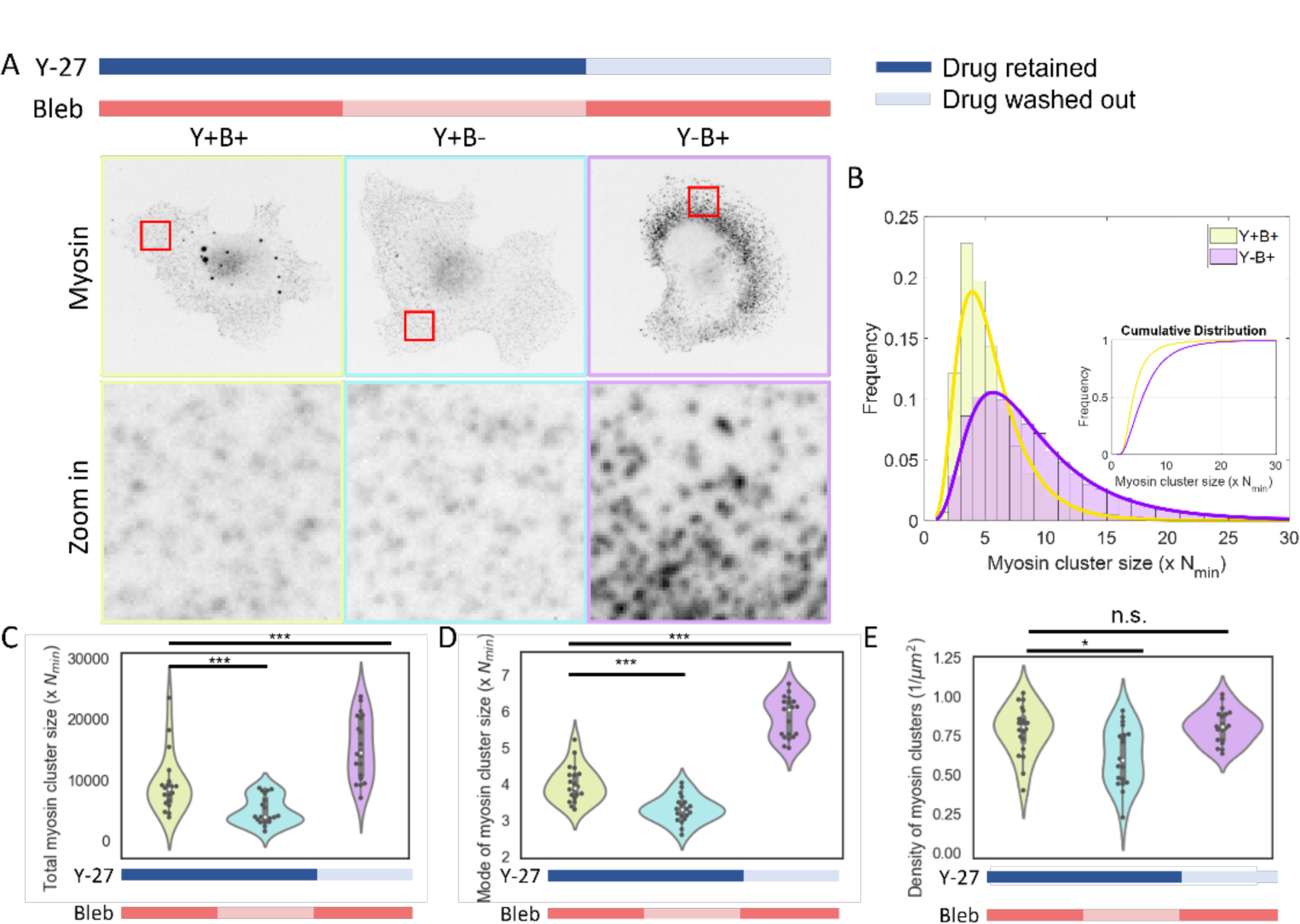
ROCK activity is sufficient for myosin clusters to grow in the absence myosin motor activity. (A) Top row shows representative myosin images of cells with or without ROCK activity and myosin motor activity. Myosin clusters were first disassembled by both Y-27632 and blebbistatin (Y+B+). Y27 or blebbistain are then removed individually (Y+B- and Y-B+). Zoomed in myosin images are shown in the bottom row. Images are inverted. Scale bar is 10 µm for the top row and 2 µm for the bottom row. (B) Histogram of myosin cluster size between Y27/blebbistatin treated and blebbistatin treated cells. Inset shows the cumulative distribution of cluster sizes. (C) The sum of myosin cluster sizes. (D) Mode of myosin cluster size distribution. (E) Density of myosin clusters. N=20 cells for each condition. Two-sample Kolmogorov-Smirnov tests were performed in (C)-(E) to test if the measured quantity differs between conditions. n.s. signifies p>0.05, * signifies p<0.05 and *** signifies p<0.001.

To decouple the role of ROCK and myosin ATPase activity on myosin cluster growth, we selectively retain one inhibitor during washout of the other. When we selectively washed out blebbistatin but retained Y-27, we see no apparent increase in myosin cluster intensity (Fig. 2A, Y+B-). The total amount of myosin contained within clusters did not shift significantly compared to the cells treated with both inhibitors (Fig. 2C). To quantify the size distribution of myosin clusters across different conditions, we fitted the myosin cluster intensity distribution to a lognormal distribution and used the mode of this distribution to represent the most probable myosin cluster size. The mode of myosin cluster intensity only marginally changed with blebbistatin washout, (Fig. 2D). This suggests that myosin ATPase activity in the absence of ROCK activity is insufficient for myosin clusters growth.

In contrast, when Y-27 is removed but blebbistatin is retained, the total amount of myosin contained within clusters across the lamella increased by ∼42% (Fig. 2A, 2C, Y-B+). From the histogram of myosin cluster sizes, we estimate that myosin clusters grow up to 4 – 15 times the size of N_min_ with ROCK activity alone (Fig. 2B, Y-B+). Compared with myosin cluster size distributions in cells treated with both inhibitors, the peak of the distribution shifts towards larger sizes and the tail widens with ROCK activity. This indicates both that the most probable myosin cluster size and the fraction of larger myosin clusters increased. This is also shown by the increasing mode of myosin cluster intensity when Y-27 is selectively washed out (Fig. 2D). Interestingly, myosin cluster density remained at similar levels in all drug conditions (Fig. 2E). This suggests that ROCK activity is sufficient for myosin clusters to grow, not through forming new clusters but through the accumulation of myosin to existing clusters.

To visualize ROCK-mediated myosin cluster growth, we conducted live-cell imaging of fluorescently-tagged myosin as Y-27 is selectively washed out in cells treated with both Y-27 and blebbistatin (Fig. 3A). As Y-27 is washed out, the intensities of myosin clusters increased over 20 minutes (Fig. 3A, B). The dynamics of individual myosin clusters reveal that most clusters are very mobile and only appear transiently for a few frames after washing out Y-27 (Fig. 3C, Supp. Movie 2). We think this does not represent rapid assembly and disassembly dynamics but instead reflects that myosin is weakly bound to F-actin due to blebbistatin treatment and, consequently, leave the field of view. It is therefore challenging to track single myosin clusters over time. Nevertheless, we observed some myosin clusters growing in intensity without apparent interaction with other clusters (Fig. 3D). This suggests that myosin clusters grow via a net association of myosin monomers or filaments to existing myosin clusters, potentiated by the increase in assembly-competent myosin driven by ROCK activity (Fig. 3E). Moreover, myosin clusters that appear at later times have higher intensities than myosin clusters that appear earlier in the washout (Fig. 3C). This suggests that myosin clusters may grow while diffusing around the cytoplasm.

**Figure 3.**
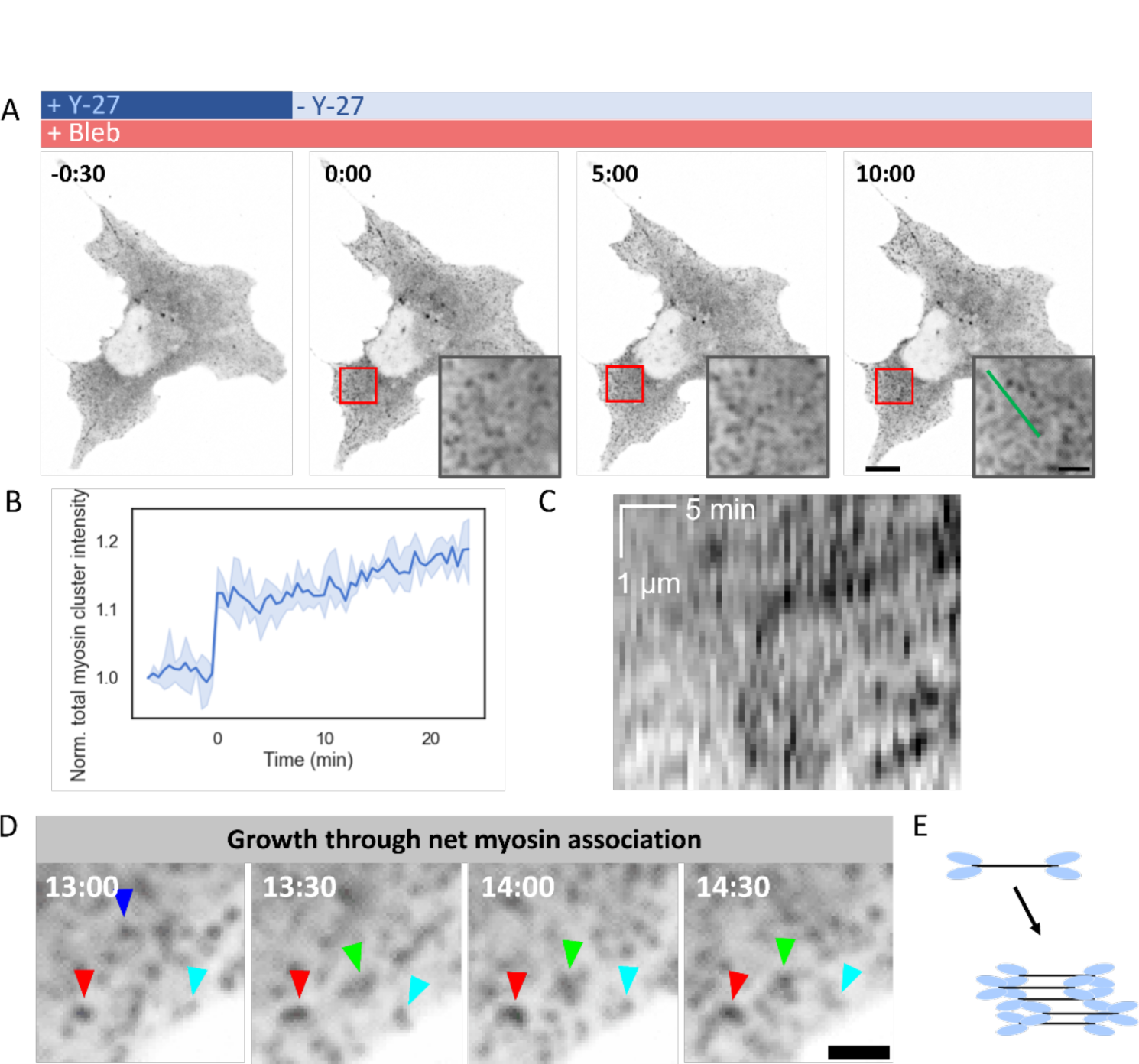
Myosin clusters grow through net association of myosin to existing clusters without myosin motor activity. (A) Snapshots of myosin clusters after washing out Y-27632 from Y-27632/blebbistatin treated cells. Time after washout is shown on the top left corner of each image. Inset shows zoom-in of myosin clusters in the red squares. Images are inverted. Scale 10 µm. Inset scale 2 µm. (B) The sum of intensities of all myosin clusters in the cell, normalized to their intensities before washout. N=3 cells. (C) Kymograph of myosin clusters along the green line in the inset in (A). Horizontal axis represents time and vertical axis represents spatial position. Time scale 5 minutes and spatial scale 1 µm. (D) Close-up snapshots of myosin cluster growth over time. Top-left indicates time after washout. Arrowheads indicate trackable myosin clusters over time. (E) Schematic of myosin association to existing clusters.

### Myosin motor activity allows myosin clusters to grow through F-actin-dependent myosin association

Our results so far show that ROCK activity is sufficient for myosin cluster growth without myosin ATPase activity. However, myosin cluster sizes are still significantly smaller than wild-type clusters (Fig. 1E & Fig. 2B). Therefore, we explored the role of myosin motor activity plays. To this end, we compared myosin cluster sizes in cells treated with blebbistatin with untreated cells (Fig. 4A). We see that the total myosin contained in clusters decreased by about 35% with blebbistatin treatment (Fig. 4C). This is consistent with previous studies showing that myosin motor activity affects myosin filament assembly (Shutova *et al*., 2012). At the level of single myosin clusters, most myosin clusters in blebbistatin-treated cells contain around 4 times more myosin than N_min_ but can contain up to 18 times more than N_min_ (Fig. 4B), which is consistent with the case when Y-27 is washed out in cells treated with Y-27 and blebbistatin (Fig. 2B, Y-B+). In contrast, while the most probable myosin cluster size only modestly increased to 5 – 6 times more than N_min_ in control cells (Fig. 4B & 4D), there is a larger fraction of myosin clusters that contain more than 15-fold of N_min_, reaching up to 28-fold of N_min_ (Fig. 4B). As in the case of ROCK-dependent myosin cluster growth, myosin cluster density remains the same with blebbistatin treatment (Fig. 4E).

**Figure 4.**
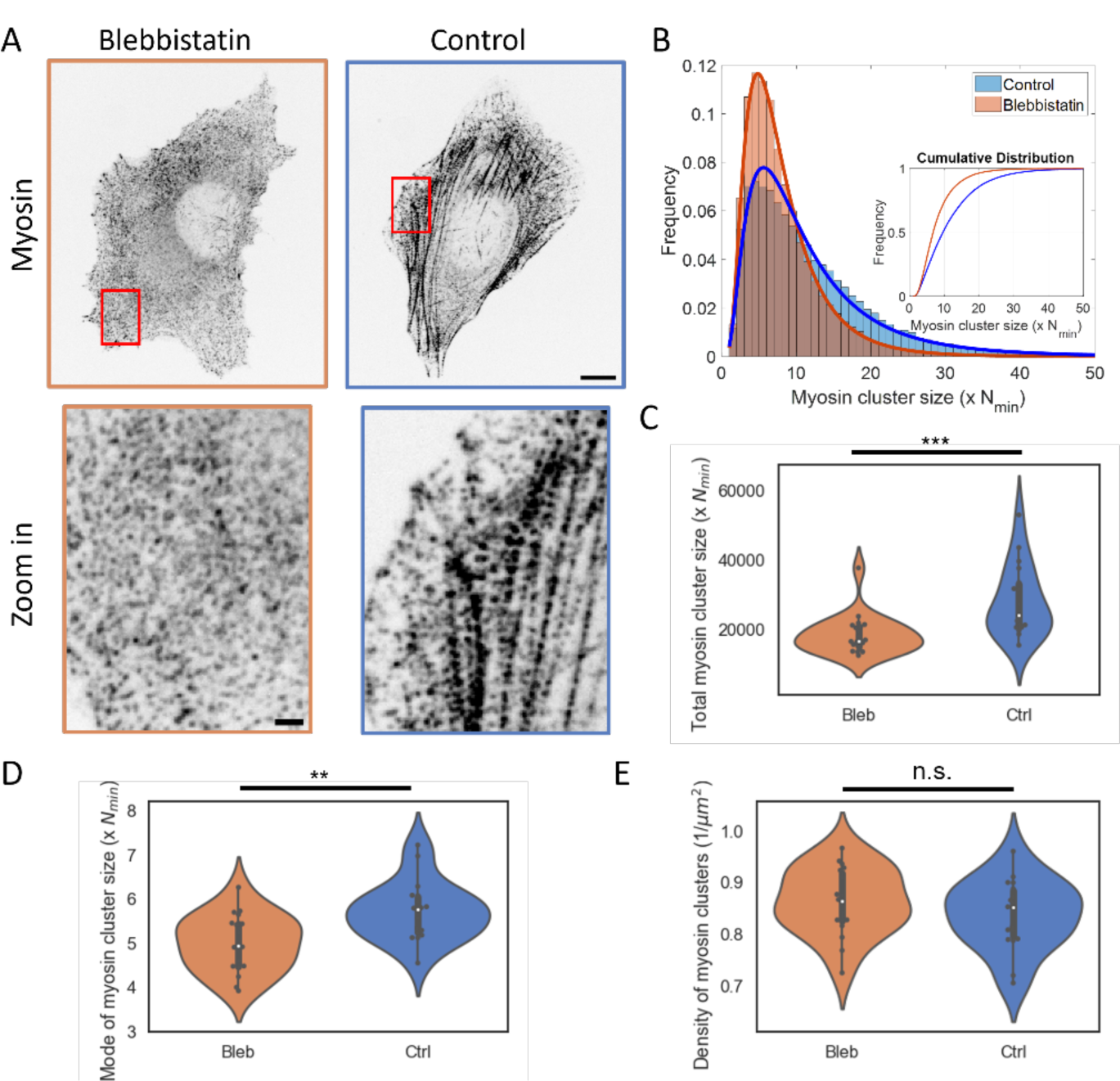
Myosin motor activity allows for myosin cluster growth. (A) Top row shows representative myosin images in cells treated with or without blebbistatin. Bottom row shows zoomed-in myosin images in the red boxes in the top row. Images are inverted. Scale bar is 10 µm for the top row and 2 µm for the bottom row. (B) Histogram of myosin cluster sizes between blebbistatin treated and control cells. Inset shows the cumulative distribution of both conditions. (C) The sum of all myosin cluster sizes. (D) Mode of myosin cluster size distribution. (E) Density of myosin clusters. N=20 cells for blebbistatin-treated cells and N=18 cells for control cells. Two-sample Kolmogorov-Smirnov tests were performed in (C)-(E) to test if the measured quantity differs between conditions. n.s. signifies p>0.05, ** signifies p<0.01 and *** signifies p<0.001.

To understand how myosin motor activity can further enhance myosin cluster size, we performed live-cell myosin imaging as we washed out blebbistatin (Fig. 5A, Supp. Movie 3). Consistent with fix and stain results, myosin cluster intensity increased as blebbistatin is washed out (Fig. 5B). During this process, myosin clusters are more stably bound, persisting for more than 10 minutes (Fig. 5C). Some myosin clusters increase intensity over time without apparent local motion or interaction with neighboring clusters, suggesting an increased net myosin association to existing clusters (Fig. 5C & D). We also observe some myosin clusters undergo local motion toward each other and merge into a brighter cluster (Fig. 5C & E), suggesting myosin cluster growth through actomyosin sliding.

**Figure 5.**
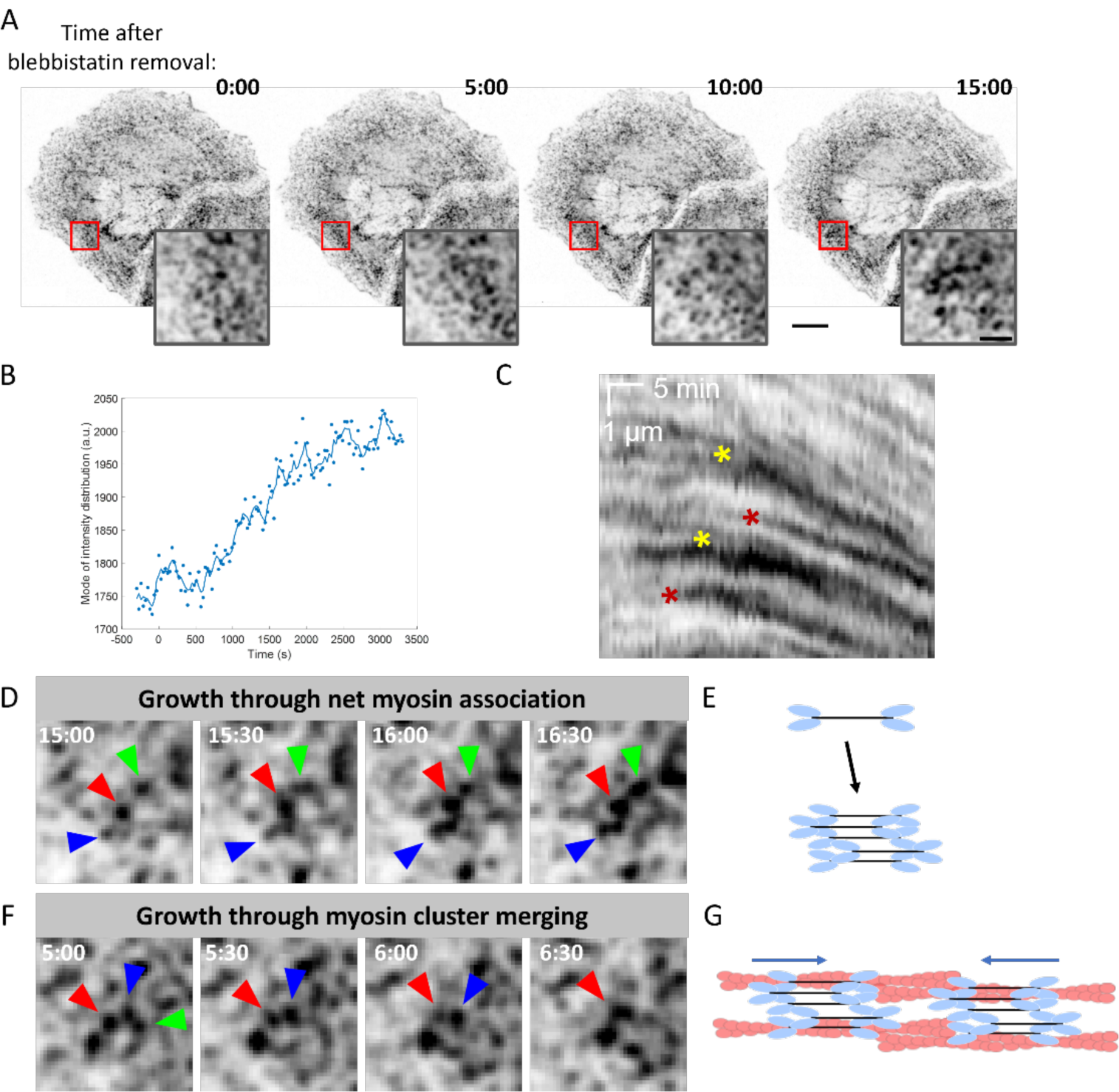
Myosin motor activity grows myosin clusters through net myosin association or cluster merging. (A) Snapshots of time-lapse imaging of myosin clusters during blebbistatin washout. Inset shows zoom-in of myosin clusters in the red squares. Images are inverted. Scale bar 10 µm. Inset scale bar 2 µm. (B) Myosin cluster intensity increases with blebbistatin washout. (C) Kymograph of myosin clusters. Maroon asterisks indicate cluster growth events through myosin association and yellow asterisks indicate two clusters merging. (D) Close-up snapshots of myosin clusters increasing intensity over time through net myosin association. Arrowheads indicate example myosin clusters. (E) Schematic of myosin association with existing clusters. (F) Close-up snapshots of myosin clusters increasing intensity through merging with another cluster driven by actomyosin sliding. Arrowheads indicate example myosin clusters. (G) Schematic of myosin cluster growth via actomyosin sliding.

Since blebbistatin does not impact RLC phosphorylation, we reasoned that myosin motor activity can only indirectly affect myosin association. Since F-actin has been proposed to regulate myosin assembly and myosin cluster growth at the leading edge (Mahajan *et al*., 1989; Fenix *et al*., 2016; Beach *et al*., 2017), we wondered if F-actin could play a role. Consistent with this idea, myosin cluster sizes across the cell positively correlate with the underlying F-actin intensity (Supp. Fig. 3). Since different stress fiber architecture contains variable amounts of myosin, we restricted our analysis to mature transverse arcs and ventral stress fibers. On these stress fibers, myosin cluster sizes correlate strongly with the intensity of F-actin bundles (Fig. 6A, C). On the other hand, there are few F-actin bundles in blebbistatin-treated cells. On these bundles, however, myosin cluster sizes showed much less positive correlation with the intensity of remaining F-actin bundles (Fig. 6B, C). Since both the slope and R^2^ of the correlation decreased with blebbistatin treatment (Fig. 6D, E), this suggests that F-actin plays a role in myosin cluster growth only in the presence of myosin motor activity and/or strong binding.

**Figure 6.**
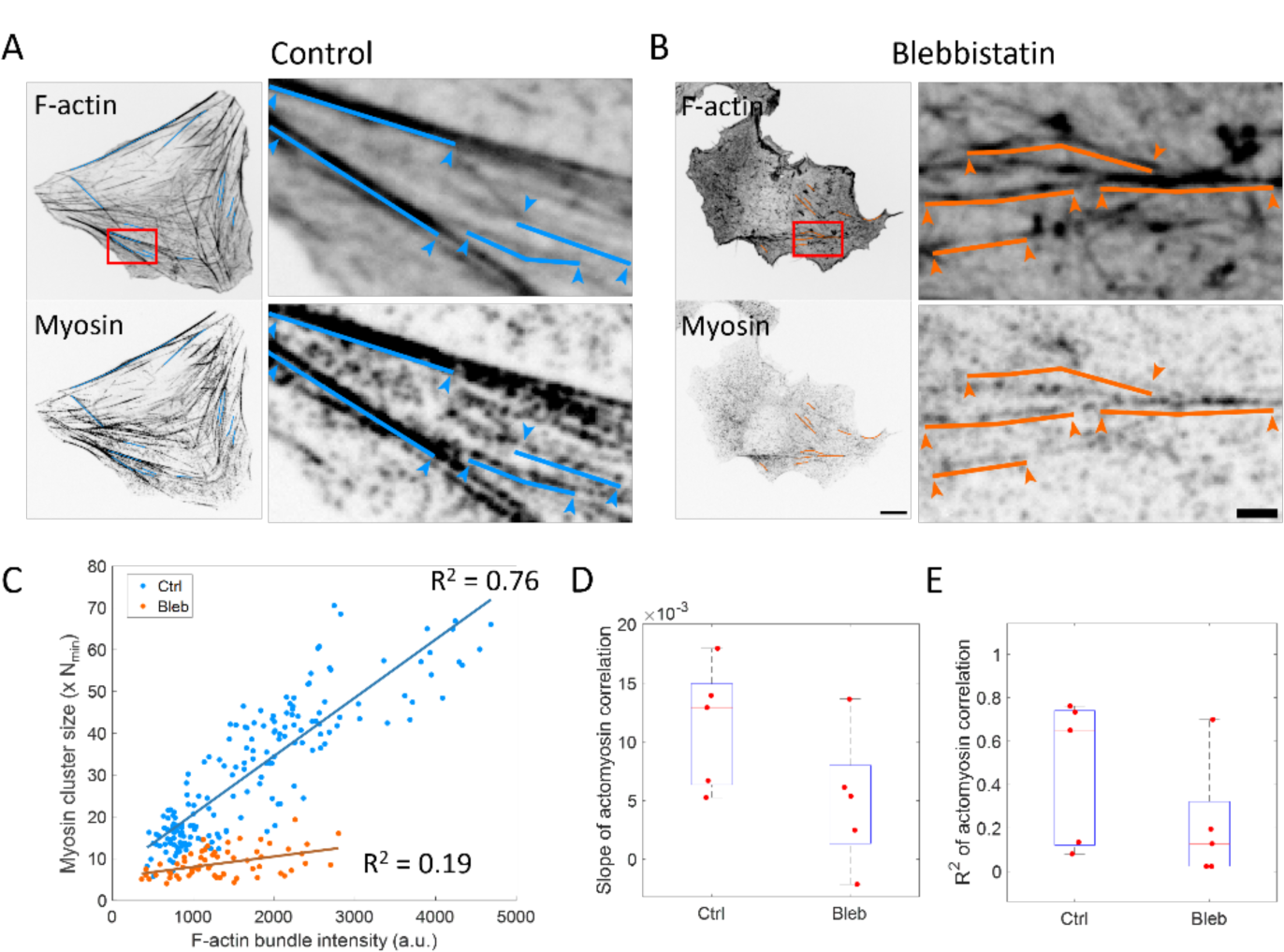
Myosin cluster sizes correlate with F-actin bundle intensity under myosin motor activity. (A) F-actin and myosin on select F-actin bundles in control cells. Left column shows all segments of F-actin bundles that are selected for analysis. Right column shows zoomed in view of a few F-actin bundles. Solid lines represent F-actin bundles where actomyosin correlation is examined. Arrowheads indicate the end points of the segment. (B) F-actin and myosin on select F-actin bundles in blebbistatin-treated cells. Scale bar is 10 µm for the whole cell image and 2 µm for the zoomed in image. (C) Correlation between myosin cluster size and F-actin bundle intensity in control and blebbistatin-treated cells. Correlation is plotted for myosin clusters on every selected bundles in a single cell. (D) Slope of the correlation. N=5 cells in each condition. (E) R^2^ of the correlation. N=5 cells in each condition.

To further understand how F-actin architecture affects myosin cluster size, we perturb the stress fiber architecture and see how myosin clusters respond. First, we disrupted F-actin crosslinking by knocking down α-actinin1 via siRNA. Under α-actinin1 knock-down, dorsal stress fibers were abolished, and transverse arc bundles reduced in actin intensity (Fig. 7A, B), which agrees with previous reports (Oakes *et al*., 2012; Feng *et al*., 2013). The total amount of myosin within clusters across the lamella does not change significantly (Supp. Fig. 4B), which is also consistent with a previous report that RLC phosphorylation levels remain unchanged (Oakes 2012). However, the sizes of myosin clusters shifted towards smaller sizes (Supp. Fig. 4A). While the most probable size of myosin clusters is similar (Supp. Fig. 4C), myosin clusters can only grow up to 20 times of N_min_ as opposed to the 30 times of N_min_ in control conditions. The density of myosin clusters also remains unchanged under α-actinin knockdown (Supp. Fig. 4D). The concurrent disruption in F-actin bundles and large myosin clusters suggests that myosin association to F-actin bundles reduced under α-actinin1 knockdown. Indeed, when we look at cluster sizes on larger transverse arcs or ventral stress fibers, myosin cluster sizes exhibit reduced positive correlation with F-actin bundle intensity (Fig. 7C). Both the slope and R^2^ of the correlation decreased with α-actinin1 knockdown (Fig. 7D & 7E). This suggests that myosin cluster sizes on F-actin bundles are reduced when F-actin bundles are disrupted.

**Figure 7.**
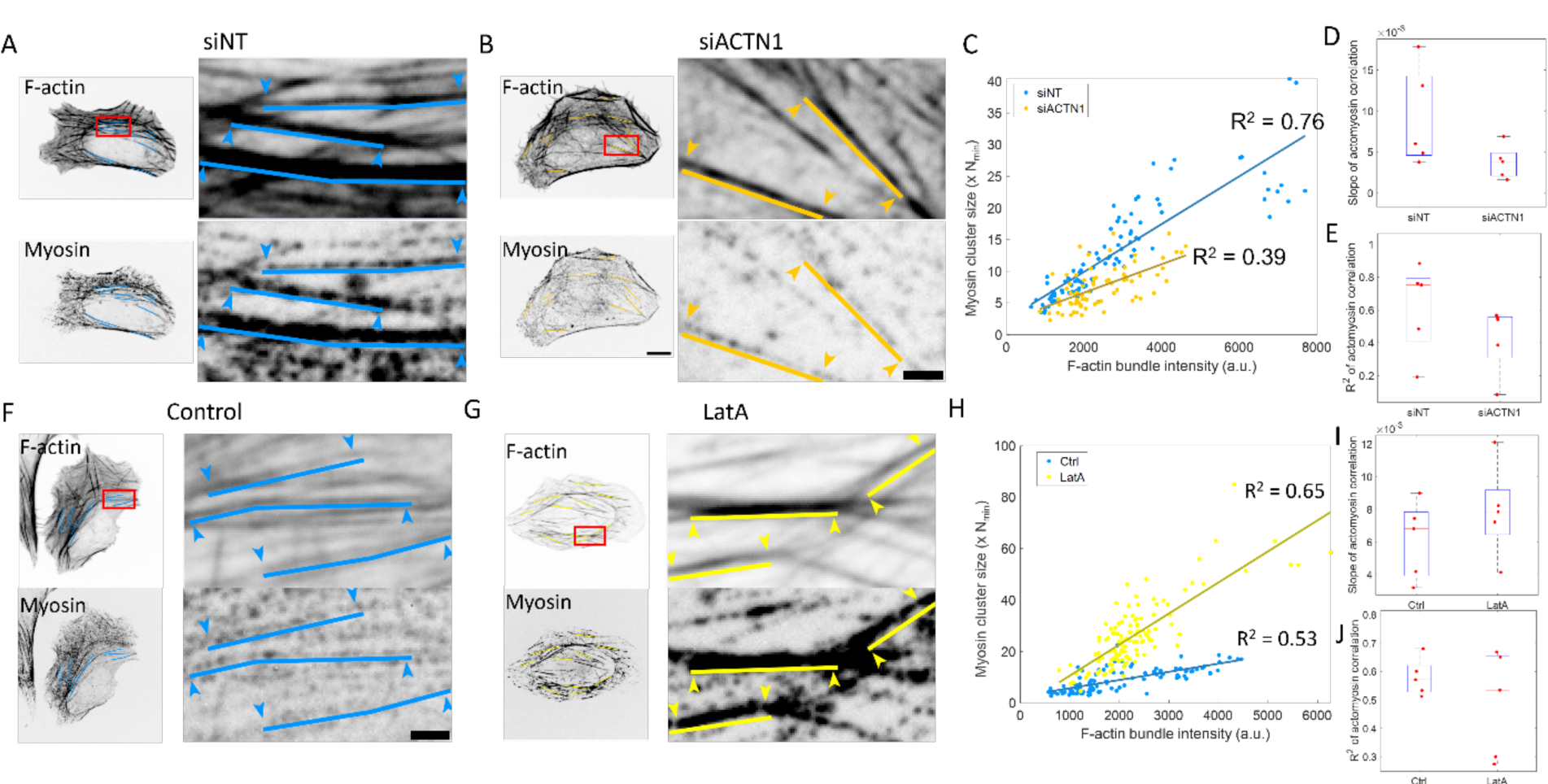
Myosin association to F-actin bundles depends on F-actin architecture. (A) F-actin and myosin on select F-actin bundles in scramble siRNA-treated cells. Left column shows all segments of F-actin bundles that are selected for analysis. Right column shows zoomed in view of a few F-actin bundles. Solid lines represent F-actin bundles where actomyosin correlation is examined. Arrowheads indicate the end points of the segment. (B) F-actin and myosin on select F-actin bundles in α-actinin1 siRNA-treated cells. Scale bar is 10 µm for the whole cell image and 2 µm for the zoomed in image. (C) Correlation between myosin cluster intensity and F-actin bundle intensity in scramble and α-actinin1 siRNA-treated cells. Correlation is plotted for myosin clusters on every selected bundles in a single cell. (D) Slope of the correlation between scramble and α-actinin1 siRNA-treated cells. N=5 cells in each condition. (E) R^2^ of the correlation between scramble and α-actinin1 siRNA-treated cells. N=5 cells in each condition. (F) F-actin and myosin on select F-actin bundles in control cells. (G) F-actin and myosin on select F-actin bundles in LatA-treated cells. Scale bar is 10 µm for the whole cell image and 2 µm for the zoomed in image. (H) Correlation between myosin cluster intensity and F-actin bundle intensity in control and LatA-treated cells. Correlation is plotted for myosin clusters on every selected bundles in a single cell. (I) Slope of the correlation between control and LatA-treated cells. N=5 cells in each condition. (J) R^2^ of the correlation between control and LatA-treated cells. N=5 cells in each condition.

On the other hand, we saw a qualitatively different result when we disrupted the smallest F-actin bundles and meshwork within the lamella by treating cells with low doses of latrunculin A (LatA). Upon treatment with 50 nM LatA, large F-actin bundles such as ventral stress fibers became the dominant F-actin architecture in cells while thin F-actin bundles were depleted (Fig. 7F, G). The sizes of myosin clusters didn’t change significantly (Supp. Fig. 4E & 4G), but the total amount of myosin within clusters decreased (Supp. Fig. 4F) due to the decrease in myosin cluster density (Supp. Fig. 4H). With thin F-actin bundles depleted, myosin clusters accumulated more on larger transverse arcs and ventral stress fibers, as seen by the larger slope of the positive correlation between myosin cluster sizes with F-actin bundle intensity (Fig. 7H, I). The R^2^ of the correlation remained similar (Fig. 7J). Together with the α-actinin1 knockdown results, our results suggest that myosin cluster sizes depend on the underlying F-actin architecture.

### Myosin cluster growth with a positive feedback is sufficient to recapitulate the broad range of myosin cluster sizes

With our experimental results showing myosin clusters growing via net association to existing clusters, we wanted to quantitatively describe myosin cluster growth. To this end, we constructed a toy model to simulate myosin cluster growth. We modeled myosin cluster growth as a Monte Carlo process, where a predetermined number of myosin filaments (N_myo_) iteratively bind randomly to a 1D grid of 10000 points. The number of grid points is fixed because myosin cluster density remained similar across almost all experimental conditions (Fig. 2E and Fig. 4E). The simulation iterates through all myosin filaments, where one myosin filament binds randomly to a grid point in each iteration. (Fig. 8A). The probability for the myosin filament on a given iteration to bind to grid point *i* is given by 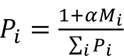, where *M_i_* is the number of myosin filaments at that grid point. The parameter α is an affinity parameter that captures scenarios where myosin filaments preferentially bind to existing myosin clusters. This is motivated by recent literature that suggests myosin filaments preferentially bind to existing myosin clusters (Fenix *et al*., 2016; Beach *et al*., 2017; Quintanilla *et al*., 2023a).

**Figure 8.**
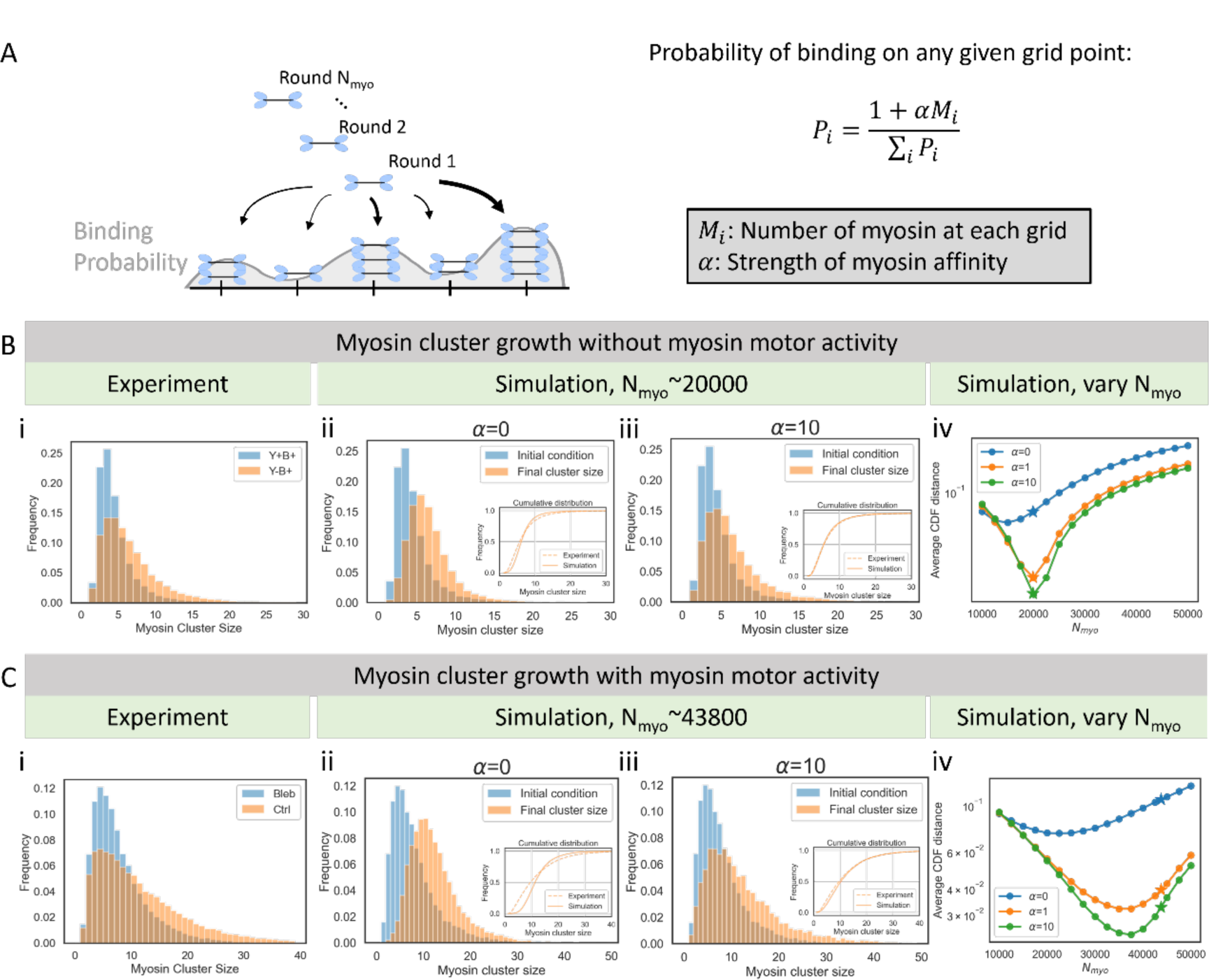
Myosin cluster growth with positive feedback is sufficient to recapitulate the broad range of myosin cluster sizes. (A) Schematic of the simulation setup. (B) Simulation of myosin cluster growth without myosin motor activity. (i) Experimentally observed myosin cluster size distribution in cells before (Y+B+) and after (Y-B+) washing out Y-27 in cells that are treated with both Y-27 and blebbistatin. (ii) Simulation when myosin binding probability is uniform across all grid points (α=0). Inset shows the cumulative distribution of the experimental and simulated results. (iii) Simulation when myosin has an affinity toward existing myosin clusters (α=10). Inset shows the cumulative distribution of the experimental and simulated results. (iv) Average distance between cumulative distributions of the experimental and simulated cluster sizes across different α and N_myo_ values. Stars show the N_myo_ determined by experimental quantifications. (C) Simulation of myosin cluster growth with myosin motor activity. (i) Experimentally observed myosin cluster size distribution in cells treated with blebbistatin or control cells. (ii) Simulation when myosin binding probability is uniform across all grid points (α=0). Inset shows the cumulative distribution of the experimental and simulated results. (iii) Simulation when myosin has an affinity toward existing myosin clusters (α=10). Inset shows the cumulative distribution of the experimental and simulated results. (iv) Average distance between cumulative distributions of the experimental and simulated cluster sizes across different α and N_myo_ values. Stars show the N_myo_ determined by experimental quantifications.

First, we use our toy model to simulate myosin cluster growth without myosin motor activity, specifically when Y-27 is washed out in cells treated with both Y-27 and blebbistatin (Fig. 2A & Fig. 8B, i). To make a meaningful comparison between the simulation and experiments, we initialized the simulation such that the size of myosin clusters on each grid matches the size distribution in cells treated with both Y-27 and blebbistatin (Fig. 8B, i, Y+B+). This is done assuming N_min_ represents one myosin filament. As a result, the initial total amount of myosin within clusters in the simulation is approximately 47000. Since our experimental results showed that ROCK-dependent myosin cluster growth increases assembled myosin by about 42% (Fig. 2C), the simulation iterates through the binding of approximately 20000 additional myosin filaments. If we assume that myosin filaments can bind to each grid point with equal probability (α = 0), corresponding to uniform growth rates across the cell, the myosin cluster sizes at the end of the simulation exhibit a Gaussian-like distribution (Fig. 8B, ii). However, this result contradicts the experimentally observed lognormal distribution when we selectively wash out Y-27 (Fig. 8B, i, Y-B+). When the affinity parameter is non-zero (α > 0), the simulated cluster sizes start to qualitatively recapitulate the lognormal distribution of myosin cluster sizes in the absence of myosin motor activity (Fig. 8B, iii). We quantified the error between simulated and experimental cluster sizes by calculating the average distance between the cumulative distributions of experimental-measured and simulated myosin cluster sizes. The error decreased with increasing α but started to converge around α = 10 (Supp. Fig. 5). This suggests that the simulated results became insensitive to changes in α, presumably due to the finite size of our simulation setup. On the other hand, the simulated results depend heavily on the number of myosin iterated through the simulation. When we scan through a range of N_myo_, the simulation error is the lowest when N_myo_ is 20000, which coincides with experimentally quantified results (Fig. 8B, iv)! This suggests that the limiting pool of available myosin is important for setting myosin cluster sizes.

We next explored if the same simulation framework and parameters can capture myosin cluster growth under myosin motor activity, specifically when blebbistatin is washed out from cells (Fig. 4A & Fig. 8C, i). To make a meaningful comparison with experiments, we initialized the simulation such that myosin cluster size on each grid matches the myosin cluster size distribution in blebbistatin-treated cells (Fig. 8C, i). Using the same assumption as before, the initial total myosin contained in clusters in the simulation is approximately 81000. Since untreated cells show 54% more assembled myosin than blebbistatin-treated cells in our experimental quantification (Fig. 4C), the simulation iterates through the binding of about 44000 additional myosin filaments. As in the case of myosin cluster growth without motor activity, uniform myosin binding probability (α = 0) fails to capture the lognormal distribution of myosin cluster sizes in control cells (Fig. 8C, ii). When the affinity parameter is non-zero (α > 0), the simulated result once again qualitatively recapitulated the experimental cluster sizes in control cells (Fig. 8C, iii). The error between simulated and experimental results decreased with increasing α but again converged around α = 10 (Supp. Fig. 5). The optimal N_myo_ that produces the least error between simulated and experimental results is around 37500, which is about 15% less than the experimentally measured N_myo_ (Fig. 8C, iv). This is likely due to the F-actin-dependent myosin cluster regulation not captured in this toy model. Taken together, our results suggest that myosin cluster sizes on stress fibers are set by a limiting pool of myosin available for cluster growth with myosin self-affinity.

## Discussion

Our results combining experiments and simulations are consistent with the picture that myosin clusters grow by myosin association to a set number of myosin clusters. We found that the most important factor that determines myosin cluster size is the limited availability of cytoplasmic myosin available to to grow clusters. We find this is regulated by both ROCK and myosin ATPase activity (Fig. 9). F-actin architecture can further regulate myosin cluster sizes (Fig. 9). On the other hand, the nucleation of myosin clusters is independent of these regulations, as we found that the density of myosin clusters remains at similar levels. This is consistent with the idea that the sizes of subcellular organelles can be controlled by the competition for a limiting pool of available components, as proposed for flagella, centrosomes, and F-actin architectures (Goehring and Hyman, 2012; Suarez and Kovar, 2016). This insight is only possible through our combination of endogenous tagging, quantitative imaging, analysis, perturbation, and modeling.

**Figure 9.**
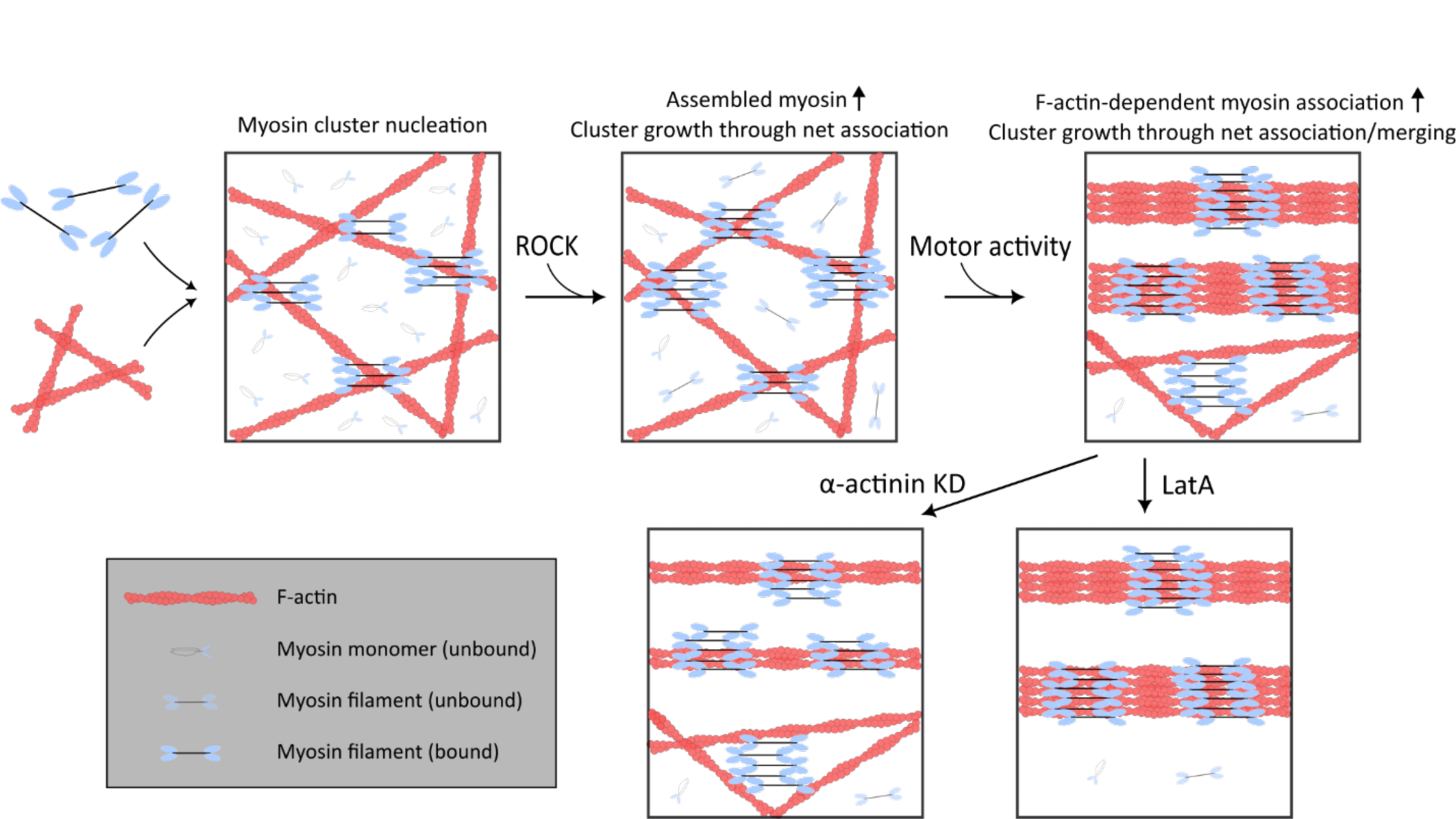
Myosin cluster sizes are set by a limiting pool of myosin that are available for cluster growth. Myosin filaments independent of ROCK activity nucleates myosin clusters. ROCK activity increases myosin assembly, which increases the pool of myosin available for cluster growth through increased net association. Myosin motor activity further enhances myosin cluster growth through F-actin-dependent myosin association. Myosin cluster sizes on F-actin bundles decreased when mature F-actin bundles are disrupted by α-actinin1 knockdown, and increased when thin F-actin bundles are depleted by LatA.

A novel finding of this study is that myosin cluster growth can occur independently of myosin motor activity. ROCK activity alone allows myosin clusters to grow from 3 times to 4-15 times the size of N_min_. Since myosin self-affinity is required for our *in silico* model to recapitulate experimental results, our results suggest that there is biochemical affinity between myosin filaments. This may explain the stack or ribbon configuration of myosin clusters observed with platinum EM (Verkhovsky *et al*., 1995; Svitkina *et al*., 1997). However, to the best of our knowledge, there is no reported molecular or electrostatic interaction between pairs of assembled myosin filaments. We speculate that the affinity can arise from numerous weak intermolecular interactions between heads, light chains, and tails of myosin monomers in neighboring myosin filaments (Yang *et al*., 2020). These interactions can be amplified when multiple myosin filaments are involved. This is supported by a previous study that reported purified NMIIA can form large clusters *in vitro* (Melli *et al*., 2018). Alternatively, the affinity between myosin filaments can be indirectly mediated by other molecular components. For example, some proteins associate strongly with myosin filaments and have been proposed to regulate myosin structures, such as myosin-18B (Jiu *et al*., 2019) or Tpm 3.1 (Meiring *et al*., 2019). This will be an interesting topic for future research to identify the cause of myosin self-affinity, and the effect on myosin clusters when this interaction is disrupted.

Myosin cluster growth can also occur with myosin motor activity mainly through the addition of available myosin to existing myosin clusters. We think that this is due to the increase in myosin assembly from myosin ATPase activity. While myosin motor activity doesn’t affect the phosphorylation of the RLC, previous studies have reported the reduction in myosin filament assembly under blebbistatin treatment (Shutova *et al*., 2012). This has been attributed to the effect of mechanical tension on myosin assembly, specifically because of the catch-bond nature of myosin-actin interaction (Veigel *et al*., 2003; Luo *et al*., 2012; Shutova *et al*., 2012). While recent studies suggest that only NMIIB exhibit catch-bond behavior while NMIIA is more like a slip-bond (Weißenbruch *et al*., 2022), NMIIA assembly can in principle be affected by NMIIB since NMIIA and NMIIB have been shown to co-polymerize into filaments (Beach *et al*., 2014; Shutova *et al*., 2014). Another possible molecular process that can result in myosin cluster growth under myosin motor activity is through the coalescence of myosin clusters resulting from their lateral motion promoted by actomyosin sliding. While coalescence occurs over a longer time scale than the association of free myosin, this can serve as a parallel mechanism for myosin cluster growth or as a means to organize the quasi-sarcomeric pattern (Thoresen *et al*., 2011).

The correlation between myosin cluster sizes and F-actin bundle size may be a result of nuanced feedback between F-actin and myosin clusters, with possible mutual regulation between myosin clusters and F-actin bundle size. On one hand, myosin can be regulated by F-actin. At the molecular level, it has been reported that myosin assembly is enhanced in the presence of F-actin (Mahajan *et al*., 1989; Applegate and Pardee, 1992; Mahajan and Pardee, 1996). Some actin-binding protein can also regulate myosin motor activity, such as tropomyosin (Barua *et al*., 2014). On the other hand, myosin can remodel the F-actin network. For example, myosin filaments can contract and remodel F-actin networks into asters (Soares E Silva *et al*., 2011; Köster *et al*., 2016; Stam *et al*., 2017). The combined effect can effectively create a positive feedback loop to give rise to the correlation between myosin cluster sizes and F-actin bundle size.

Our understanding of myosin cluster regulation is consistent with the idea that the number and size of subcellular organelles are tightly controlled (Marshall, 2016; Chen and Levy, 2022; Amiri *et al*., 2023). One model of organelle size control is that a limited pool of cytoplasmic components sets the size of these structures. This concept has been demonstrated in several organelles, such as the flagella (Ludington *et al*., 2012), centrosomes (Decker *et al*., 2011), and mitotic spindles (Mitchison *et al*., 2015). Similarly, structures that depend on the same building blocks can compete for these limited components, as observed with different F-actin architectures (Suarez and Kovar, 2016; Kadzik *et al*., 2020) and myosin clusters on various actomyosin structures (Beach *et al*., 2017; Najafabadi *et al*., 2022). Our paper contributes to this idea by quantitatively demonstrating its applicability in stress fibers of non-muscle cells. We believe that our approach, combined with advanced microscopy, molecular biology, and perturbation techniques, will lead to further insight into how cells balance and control organelles.

## Material and methods

### Cell culture

U2OS cells were cultured in McCoy’s 5A Medium (Sigma-Aldrich) supplemented with 10% FBS (Corning) and 2 mM L-glutamine (Invitrogen). To visualize myosin, mScarlet was knocked in at the C-terminal locus of the MYH9 gene using CRISPR. The target/Cas9 plasmid pSpCas9(BB)-2A-Puro (PX459) V2.0 (http://www.addgene.org/62988/) with target sequence AGGTAGATGGCAAAGCGGAT was engineered according to established protocols (Ran et al, 2013). To create a donor plasmid, pUC57 was digested with EcoRI and StuI and purified. A four-piece Gibson assembly was then performed. Three gBlocks (5’ HDR, linker/mScarlet, and 3’HDR) were obtained from IDT with overlapping extensions to mediate Gibson assembly. The 5’ HDR arm is 867 bp of genomic sequence terminating immediately prior to the endogenous STOP codon. Silent mutations (AaGTtGAcGGaAAAGCGGATGGt) were placed in the target sequence to prevent Cas9 binding and cleavage of the donor plasmid. The linker/mScarlet was placed in-frame with the coding sequence and consists of an 18 amino acid GS rich linker and mScarlet-I fluorophore. The 3’ HDR arm is 684 bp immediately downstream of the MYH9 coding sequence. U2OS cells were transfected with both Cas9 and donor plasmids. After ∼ 1 week in culture, a polyclonal mScarlet positive population was obtained using fluorescent activated cell sorting (FACS).

For fluorescent-recovery after photobleaching (FRAP) experiments, eGFP was knocked in at the N-terminal locus of the MYH9 gene instead. The following modifications were made for the CRISPR procedure. The target sequence for the guide RNA is AAACTTCATCAACAATCCGC. Donor plasmid was generated in pUCIDT-AmpR with internal mEGFP flanked by 5’ and 3’ HDR. The 5’HDR is 498 bp upstream of endogenous start and 3’HDR is 383 bp downstream of endogenous start. All cells were previously treated with BM cyclin (Roche) per manufacturer instructions to eliminate mycoplasma.

### Drug Perturbations and Gene Knockdown

Cells were treated with 40 µM Y-27632 (EMD Millipore) or 50 µM blebbistatin (MilliporeSigma) for at least 30 minutes. Latrunculin A (Sigma-Aldrich) is used at a sub-saturating concentration of 50 nM for 30 minutes. To knock down alpha-actinin, ON-TARGETplus SMARTpool human ACTN1 siRNA is used per manufacturer instructions (Horizon Discovery). For siRNA control, ON-TARGETplus non-targeting siRNA is used (Horizon Discovery).

### Immunofluorescence

Cells were first washed with warm PBS. Then cells were fixed and permeabilized with 4% paraformaldehyde (EMS) and 0.5% Triton X-100 (Fisher Scientific), diluted in 1.5% BSA (Fisher Scientific) and cytoskeletal buffer (0.01M MES, 0.003M MgCl_2_, 0.138M KCl, 0.002M EGTA, pH 6.8). To visualize F-actin, cells were then washed with PBS then incubated with Alexa-647 phalloidin (Invitrogen) at 1:1000 dilution, 0.5% Triton X-100, and 1.5% BSA diluted in cytoskeletal buffer. Cells were then washed with PBS and mounted onto a coverslip with ProLong gold antifade reagent (Invitrogen).

### Microscopy and Live Cell Imaging

Cells were imaged on an inverted Nikon Ti-E (Nikon, Melville, NY) with a Yokogawa CSU-X confocal scanning head (Yokogawa Electric, Tokyo, Japan) and laser merge model with 491, 561, and 642nm laser lines (Spectral Applied Research, Ontario, Canada). Images were collected on a Zyla 4.2 sCMOS Camera (Andor, Belfast, UK). A 60x 1.2 Plan Apo water (Nikon) objective was used to collect images. MetaMorph Automation and Image Analysis Software (Molecular Devices, Sunnvyale,CA) controlled all hardware. For quantitative myosin cluster size analysis, all imaging conditions are controlled to be the same in each set of experiments, including laser intensity and exposure time. For fluorescence recovery after photobleaching (FRAP) experiments, Airyscan imaging was performed on a Zeiss LSM 980 microscope equipped with the Airyscan 2 detector. Images were acquired using the MPLX SR-4X mode and processed by Zen Blue 3.0 software using the Airyscan processing feature with default settings. For live cell imaging, cells were mounted on an imaging chamber (Chamlide) and maintained at 37 °C. For live cell imaging, cell medium was replaced with Dulbecco’s Modified Eagle Medium without phenol red (Corning) supplemented with 10% FBS, 2mM L-glutamine, 1% penicillin-streptomycin (Corning), 10mM HEPES (Corning) and 30 µL/mL Oxyrase (Oxyrase Inc.).

### Image Processing

For the quantification of myosin cluster sizes, confocal images were captured ∼ 2 µm around the focal plane to capture all intensities and z projection ∼ 1 µm was performed around the focal plane where myosin is in focus. To remove diffuse myosin background, immunofluorescence images were first background subtracted using a top-hat filter with a circular element with a diameter of 50 pixels. Then myosin clusters were localized using a feature-finding code (Crocker and Grier, 1996). The identified features were filtered based on the general criteria: minimum intensity, maximum radius of gyration, maximum eccentricity, and intensity divided by radius of gyration. Features outside of the cell area and features closer than 2 pixels apart (0.2 microns) were also excluded. Myosin puncta intensity were then fitted to a lognormal distribution using maximum-likelihood estimation (Matlab).

Live-cell imaging movies were first corrected for photobleaching. The photobleaching curve was generated by summing all pixel intensities within a cell. The photobleaching curve is then fitted empirically to a double exponential decay function of the form 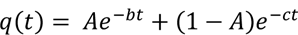. Then intensities were corrected by dividing the imaging intensity to the curve 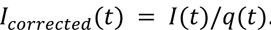. Afterwards, movies were either denoised by applying a spatial Gaussian filter (sigma=1 pixel) or by Noise2Void (Krull *et al*., 2019).

To analyze FRAP experiments, both FRAP and control regions were identified. A double normalization scheme is used to normalize data and correct for photobleaching 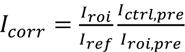. The FRAP curve is fitted to *I*(*t*) = *C*(1 – *e*^-α*t*^) to extract the recovery half time ϊ_1/2_ = *ln*2/α and the mobile fraction *C*. All analysis codes are available upon request.

### Simulation of Myosin Cluster Growth

The simulation setup is summarized in the main text. To match the initial myosin cluster size distribution in the simulation with experimental conditions, each grid point is initialized with a cluster size that is randomly sampled from the experimental cumulative distribution of cluster sizes. After all grid points have been initialized, the initial total cluster size can be calculated, and N_myo_ can be determined by percent increase of total myosin cluster sizes in experimental quantifications as described in the main text. The simulation iterates through the random binding of N_myo_ myosin filaments to a grid. In each iteration, one myosin filament binds to a grid chosen by the default Python function *choices*, with the probability to choose each grid being 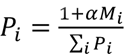 as described in the main text. Code is available on Github.

## Acknowledgement

The authors thank Melissa Quintanilla, Steven Redford and Ed Munro for insightful discussions. This work is supported by NIH RO1 GM104032.

---

**Supplementary Fig. 1.**
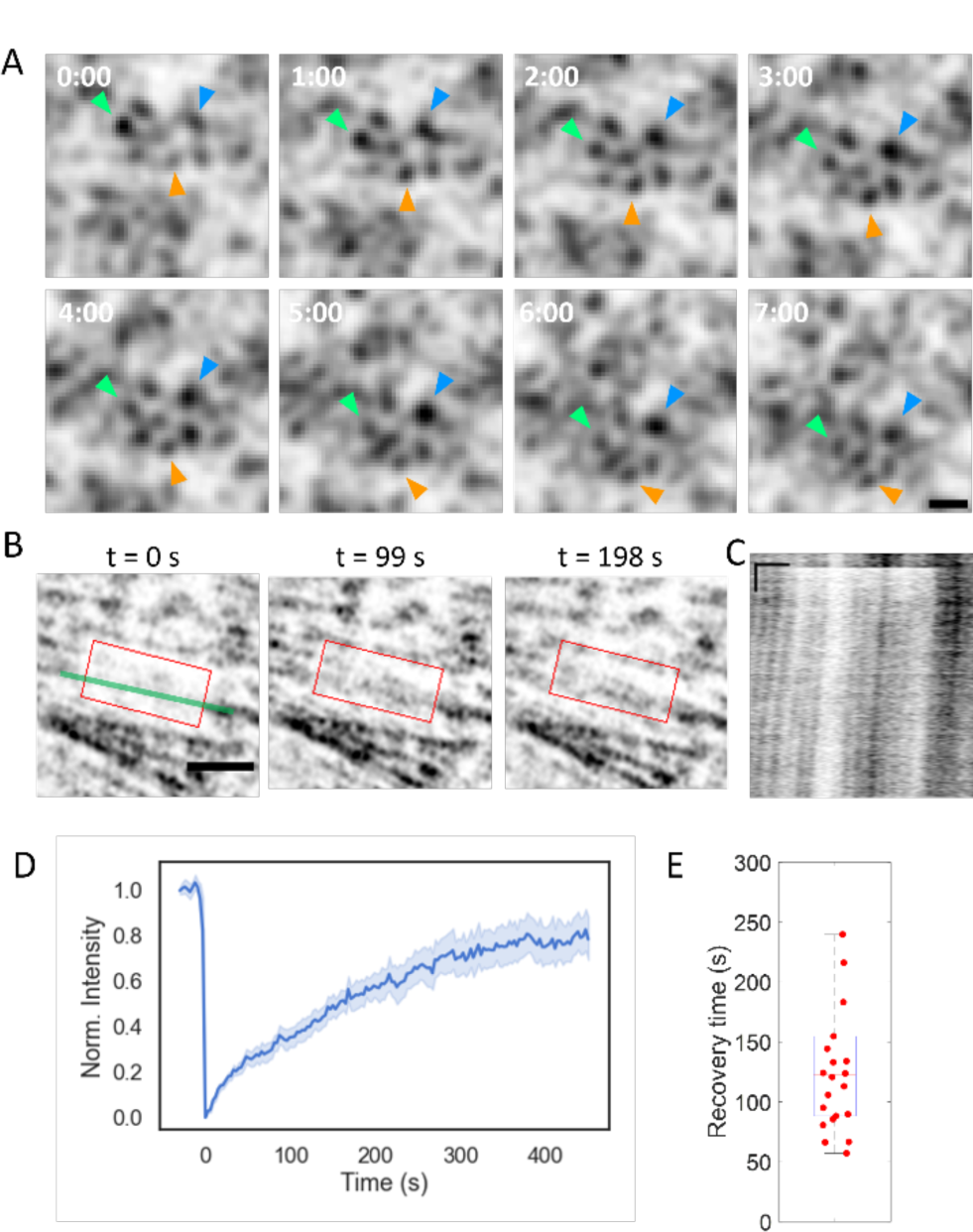
Myosin cluster dynamics. (A) Snapshots of timelapse myosin cluster imaging. Arrow heads track the same myosin cluster over time. Time stamps shown on the upper left of the images (minutes:seconds). Scale bar 2 µm. (B) Representative images of fluorescence recovery after photobleaching (FRAP) experiments. Time represents time after photobleaching. Red box indicates photobleached region. Scale bar 2 µm. (C) Kymograph along a photobleached stress fiber. The stress fiber is indicated as a green line in (B). Horizontal axis indicates spatial position with a scale bar of 1 µm. Vertical axis indicates time with a scale bar of 1 minute. (D) FRAP recovery curve. N = 22 ROIs in 6 cells. (E) FRAP recovery half time.

**Supplementary Fig. 2.**
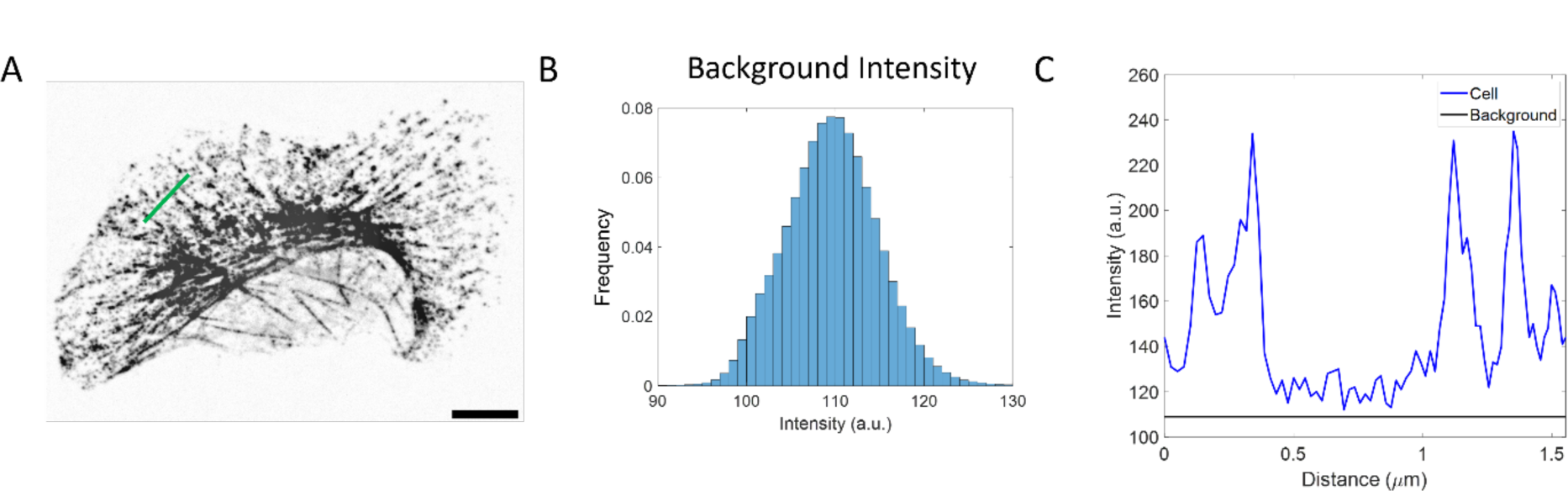
Myosin background intensity. (A) Representative image of endogenously-tagged myosin after fixation and permeabilization. (B) Histogram of background pixel intensities outside of the cell. (C) Fluorescence intensities along the green line in (A). Black horizontal line at 110 a.u. indicates the mean of background pixel intensity.

**Supplementary Fig. 3.**
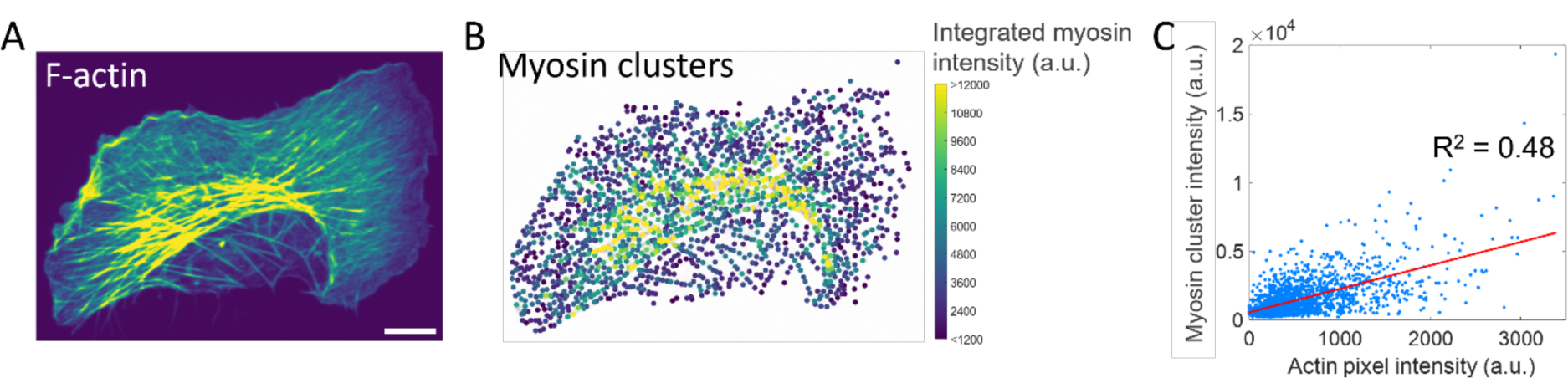
Myosin cluster intensity correlates with F-actin intensity. (A) Representative F-actin image shown on the same color bar as myosin clusters. (B) Size of myosin clusters shown color-coded by the integrated intensity of myosin puncta. (C) Correlation between myosin cluster intensity and F-actin pixel intensity across the whole cell.

**Supplementary Fig. 4.**
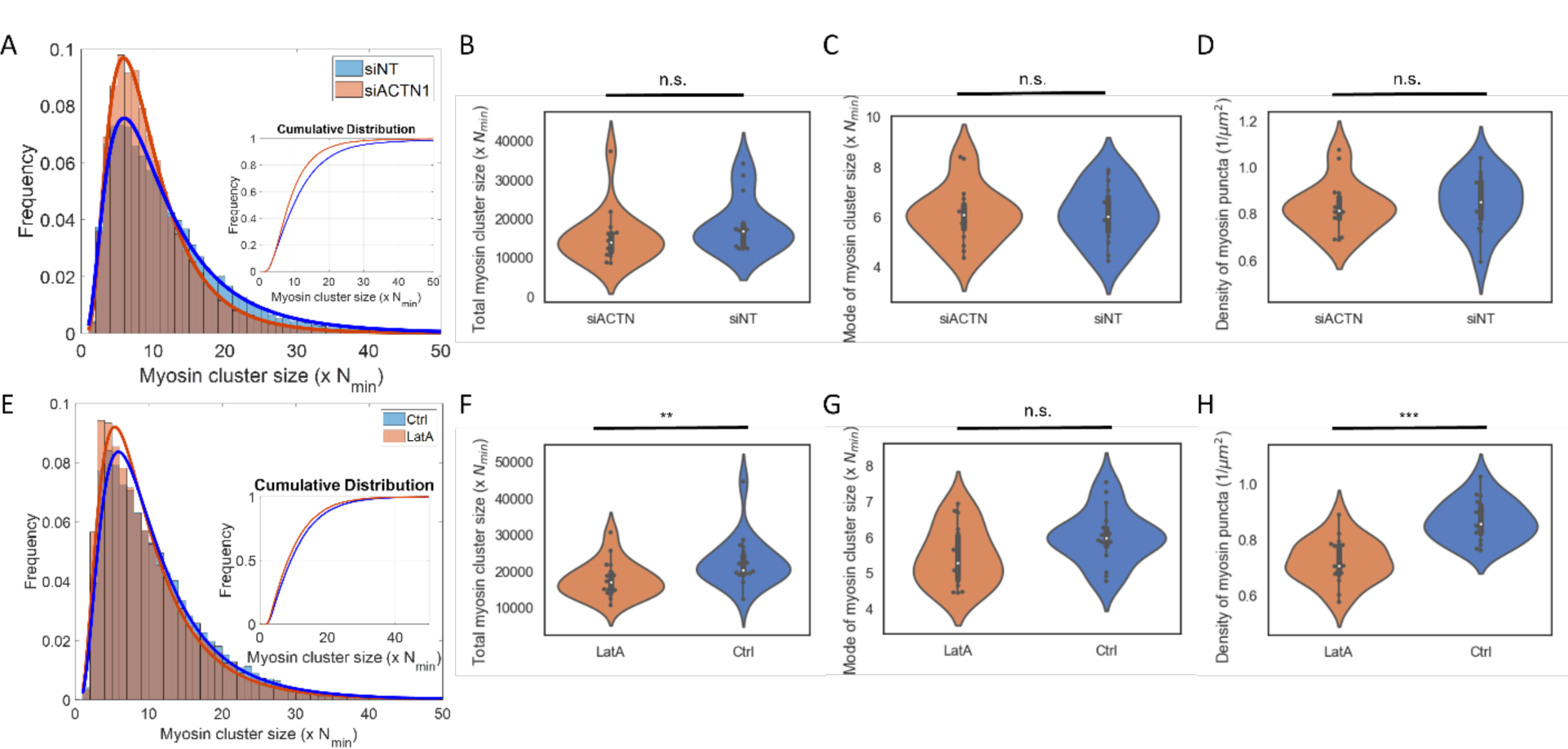
Myosin cluster sizes under F-actin perturbation. (A) Histogram of myosin cluster sizes of cells transfected with scramble siRNA and ACTN1-siRNA. Inset shows the cumulative distribution of cluster sizes. (B) Total myosin cluster sizes between cells transfected with scramble siRNA and ACTN1-siRNA. (C) Mode of myosin cluster size distribution between cells transfected with scramble siRNA and ACTN1-siRNA. (D) Density of myosin clusters between cells transfected with scramble siRNA and ACTN1-siRNA. (E) Histogram of myosin cluster sizes in control and LatA-treated cells. Inset shows the cumulative distribution of cluster sizes. (F) Total myosin cluster size in control and LatA-treated cells. (G) Mode of myosin cluster size distribution in control and LatA-treated cells. (H) Density of myosin clusters in control and LatA-treated cells. Two-sample Kolmogorov-Smirnov tests were performed in (B)-(D) and (F)-(H) to test if the measured quantity differs between conditions. n.s. signifies p>0.05, ** signifies p<0.01 and *** signifies p<0.001.

**Supplementary Fig. 5.**
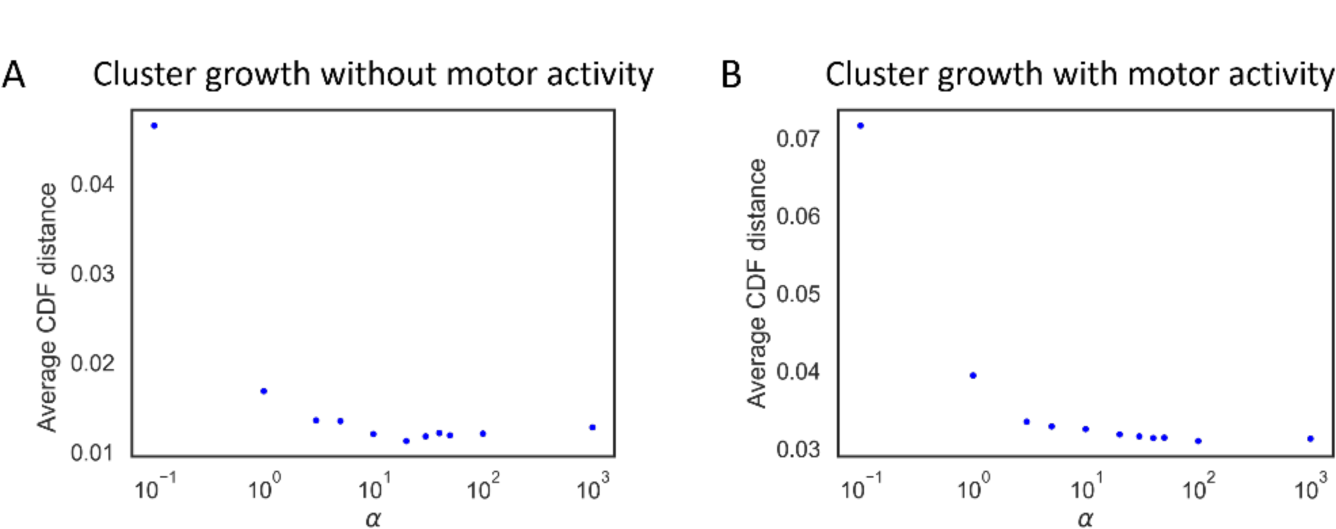
Simulation of myosin cluster growth recapitulates experimental cluster sizes with myosin self-affinity. (A) Simulation for myosin cluster growth without myosin motor activity. Difference between simulated and experimental cluster sizes are represented by the average distance between their cumulative distributions. Larger α indicates larger myosin self-affinity. (B) Simulation for myosin cluster growth with myosin motor activity.

## Supplemental Movies

Supplementary Movie 1 Endogenous myosin imaging in U2OS cells. Color is inverted. Time stamp indicates minutes and seconds.

Supplementary Movie 2 Myosin imaging in cells when Y-27 is washed out from Y-27 and blebbistatin-treated cells. Color is inverted. Time stamp indicates minutes and seconds. Images are denoised with Noise2Void.

Supplementary Movie 3 Myosin imaging in cells when blebbistatin is washed out from cells. Color is inverted. Time stamp indicates minutes and seconds.

